# Chemical interactions in polyethylene glycol-induced condensates lead to an anomalous FRET response from a flexible linker-fluorescent protein crowding sensor

**DOI:** 10.64898/2026.02.16.706251

**Authors:** Anshuman Mohapatra, Jeanpun Antarasen, Danielle R. Latham, Marisa A. Barilla, Caitlin M. Davis, Lydia Kisley

## Abstract

The cellular cytosol is a crowded environment. Biomolecular Förster resonance energy transfer (FRET) sensors have been developed to measure crowding in cytosol mimics comprised of synthetic polymers such as polyethylene glycol (PEG) and Ficoll that impart an excluded volume effect. In the current study, we explore the unsolicited role of PEG in driving the phase separation of a protein crowding sensor, AcGFP1/mCherry-FRET crowding helix 2 (CrH2), into fluorescent puncta. In contrast, a DNA-based crowding sensor (CrD), with an Alexa488/Cy5 FRET pair, does not form puncta under the same crowding conditions. Using fluorescence recovery after photobleaching imaging, we uncover the liquid-like physical properties of the PEG-induced puncta. Two-color fluorescence microscopy imaging reveals crowder-induced inhomogeneity, concentration variations, and partition coefficient across the dilute and dense phases of the liquid puncta, which remain largely underexplored in bulk fluorometry measurements. Thus, the average crowding sensor response may originate from an aqueous biphasic system, reporting an erroneous average response instead of distinct levels of crowdedness. A comparison of excluded volume effects conferred by Ficoll and PEGs of various molecular weight ranges shows the influence of size, concentration, excluded volume, and chemical composition on the CrH2 sensor response. We demonstrate that PEGs enable phase separation and alter sensor response through a mechanism that may be driven by polymer interactions with the flexible hinge region of CrH2. Overall, we determine the biophysical mechanisms underlying PEG-induced condensation of CrH2 and demonstrate a CrD sensor as an alternative that does not undergo phase separation.

## Introduction

Cells have a significant portion of their volume occupied by macromolecules, with concentrations ranging from 50 to 500 mg/mL.^1–4^ A change in macromolecular content enables the regulation of up to 30% of the cell volume during mitosis or in response to stress conditions.^5^ Moreover, macromolecular crowding enables, regulates, and impacts the molecular interactions, assembly, and dynamics in living cells.^6–8^ The participation of proteins and nucleic acids in crowding phenomena enables complex formation, facilitating cellular storage,^9^ stress response,^10^ and signal transduction^11^ in various cellular pathways.^12^ Under crowded conditions, these macromolecules selectively undergo conformational transitions to attain energetically favorable states.^13–15^ Crowding alters macromolecules’ rotational and translational degrees of freedom,^16^ leading to folding,^14^ association,^17^ aggregation,^17^ and excluded volume effects^15,18^ that can influence supramolecular complex formation.^19^

Molecular crowding and excluded volume effects, initially introduced by Allen Minton,^15^ have been estimated by calculating macromolecular volume fractions using both direct^1^ and indirect approaches.^20,21^ Volume changes are typically observed via osmotic upshifts,^19^ protein content,^22^ and global diffusion coefficient estimation.^23,24^ Yet, spatiotemporally, the cell is constantly changing in organization.

Förster resonance energy transfer (FRET)-based sensors with protein or DNA backbones have been developed and applied to spatiotemporal crowding measurements in synthetic and cellular microenvironments.^12,19,20,25–27^ These crowding sensors are typically designed with a FRET pair connected through a conformationally flexible hinge framework.^26^ The hinge serves as a fulcrum, allowing crowding-induced and compaction-mediated reductions in the Förster distance between the donor and acceptor pairs.^26^ Photophysically, the donor fluorescence quenching via non-radiative energy transfer enables ratio-metric acceptor fluorescence enhancement in response to crowding.^26^

FRET-based protein crowding sensors have been used in a wide range of concentrations and environments that can lead to complications in the fluorescence readout. It is generally considered safe to use the sensor at concentrations below 200 nM to avoid intermolecular FRET.^26–28^ *In vitro* studies have used crowding sensors at concentrations up to 5 µM.^12,20,29–32^ However, quantification of sensor overexpression in bacteria^26^ and transient cellular^27,33^ overexpression is often not reported, which makes it challenging to understand the amount of sensor present in cell measurements.

In vitro, crowding is typically achieved using molecular crowders such as bovine serum albumin (BSA), polyethylene glycol (PEG), dextran, and Ficoll.^26,34^ The synthetic polymer PEG is considered relatively inert and is used to approximate the intracellular concentration of macromolecules.^26,31,34^ However, PEG is also known to reduce the entropic barrier, enabling crowding-induced phase separation of proteins.^34,35^ The phase properties of PEG are facilitated by co-condensation,^34^ increased dehydration entropy,^36^ osmotic pressure effects,^35^ and depletion forces.^35,37^ In addition to the widely known disordered proteins that undergo phase separation, globular and well-folded proteins have also been observed to undergo the same fate.^38–40^ FRET-based protein crowding sensors are no exception to phase separation.^26^ Moreover, crowding sensors are known to have artifacts, including but not limited to maturation,^25^ stability,^20^ and coacervate formation.^26^ Understanding the nature of the crowding sensor is essential when reporting under higher osmotic levels,^19^ different cell cycle stages,^19,41^ chemical agents,^42^ and crowders.^43^

The current study investigates the anomalous FRET response from biomolecular crowding sensors in the presence of synthetic crowders that cause phase separation. We observe that the AcGFP1/mCherry-FRET crowding helix 2 (CrH2) crowding sensor phase separates in the presence of PEG but not Ficoll.^29^ Two-color microscopy imaging reveals CrH2 forming fluorescent punctate structures in the presence of PEG.^26^ In parallel, we compare the crowder-induced FRET response of CrH2 with a DNA-based crowding sensor “Crowding DNA helix” (CrD), which, in contrast, does not form puncta. The CrH2 that undergoes PEG-induced phase separation exhibits altered diffusion, viscoelastic, biophysical, and photophysical properties.^32^ The inhomogeneity via phase separation impacts *in vitro* sensing and calibration, which remain deemphasized in earlier reports.^26^ Using fluorescence recovery after photobleaching (FRAP), we have uncovered the liquid-like properties of CrH2 within the PEG-induced fluorescent puncta, confirming phase separation and condensate formation. We investigate the aqueous biphasic system using partition coefficient analysis of microscopy images to gain a better understanding of the average sensor response. The excluded volume effects conferred by the synthetic crowder Ficoll PM 70 (70 kDa) and PEGs of 1 – 35 kDa molecular weight on the crowding sensor were quantified to understand the interplay between size, concentration, excluded volume, and crowder chemistry sensor phase and response.

## Materials and Methods

### Materials

The respective vendors supplied the following materials: mCherry (Abcam #ab199750), eGFP (Abcam #ab51992), HEPES (Thermo Scientific #BP310), and Ficoll PM 70 (Cytiva #GE17-0310). PEG 1 kDa (#81189), PEG 8 kDa (#89510), PEG 20 kDa (#95172-250G-F), PEG 35 kDa (#94646-250G-F), Tris EDTA-Buffer solution (93283), Trolox (#648471), and nucleus-free water (#W4502) were all obtained from Sigma Aldrich. Dithiothreitol (DTT) (#DTT25) was from Goldbiotech. H_2_O_2_ was 30% Certified ACS grade from Thermo Scientific, and NH_4_OH was Certified ACS Plus grade from Fisher Chemical. All water used for analysis was purified from an ELGA system to Type I purity with resistivity ≥ 18.2 MΩ unless noted otherwise.

### Expression and Purification of Protein

The CrH2 protein (Table 1) was expressed and purified as described in earlier work by Davis and co-workers.^29,32^ Briefly, a pDream2.1 plasmid containing CrH2 was transformed into BL21-CodonPlus(DE3)-RIPL *E. coli* (Agilent). The transformed cells were subsequently grown in Lennox LB Broth (Fisher Scientific) at 37 °C until they reached an OD_600_ of 0.6. The cells were then induced with 1 mM isopropyl-β-D-thiogalactopyranoside (Inalco) and grown for 16 hours at 20 °C. The cells were harvested using centrifugation, resuspended in lysis buffer, and lysed via sonication. After sonication, the lysate was clarified by centrifuging and filtering, then loaded onto a Ni-NTA column for purification. The purified protein was confirmed using SDS-PAGE and then dialyzed into 20 mM sodium phosphate buffer, pH 7. The purified protein was stored at −80 °C. Unless otherwise stated, the experiments were performed within one week of freshly thawing (post-thaw protein stocks were kept at 4 °C) a 10 µM aliquot of the protein without further downstream processing.

**Table 1:**
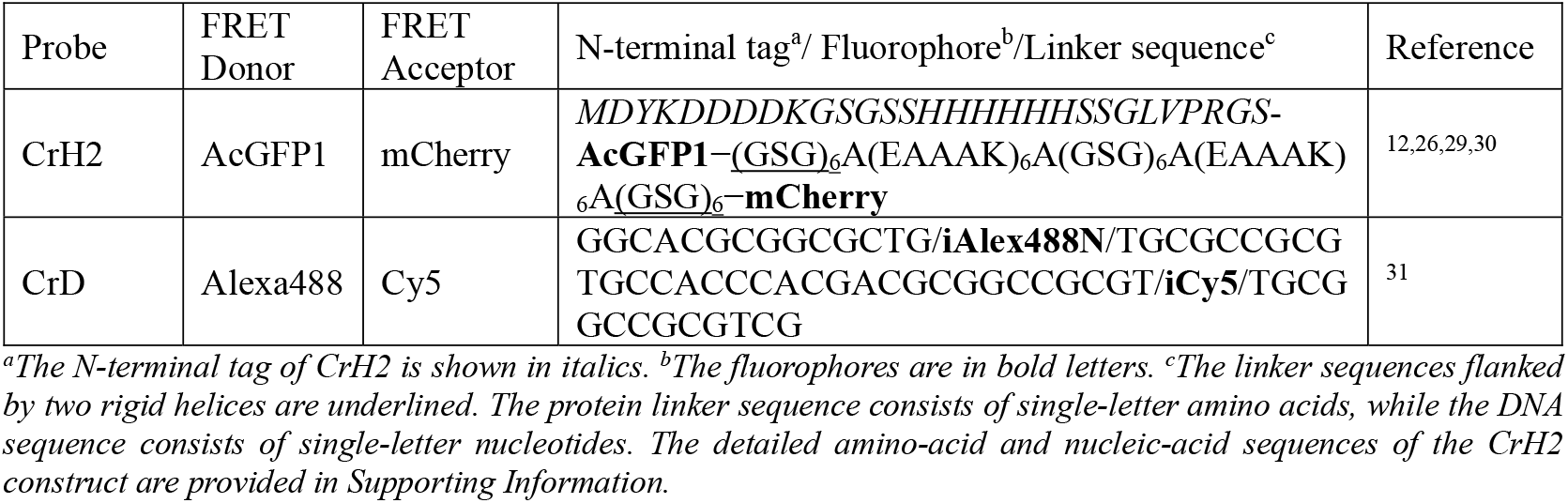
Sequences and Fluorescent Labels of CrH2 and CrD Crowding Sensors.

### Ultracentrifugation of the Protein Sample

To remove any preformed aggregates, a freshly thawed batch of CrH2 was ultracentrifuged before two-color fluorescence microscopy using an OptimaTM MAX-XP Ultracentrifuge at 100,000 x g with a TLA-120.1 rotor for 30 minutes at 4 °C, using 500 µL Beckman centrifuge tubes (#343776). The supernatant was recovered, and SDS-PAGE (Figure S1) and UV/Vis spectra (Figure S2) were recorded to ascertain purity and concentration, respectively. The UV/Vis spectra were recorded using a JASCO V-750 spectrophotometer from 250 to 700 nm at a scan rate of 100 nm/min using a 1 cm ultra-micro Hellma fluorescence quartz cuvette at 25°C. Extinction coefficients of the AcGFP1 and mCherry were used for the concentration estimations (Table S1).

### Preparation of Crowding Agents

The Ficoll solution was prepared by adding 4 g of Ficoll PM 70 powder to 5 mL of water and incubating it at 55 °C in a water bath overnight without agitation. In the morning, a clear solution of Ficoll PM 70 at ∼500 mg/mL was obtained and briefly sonicated in a water bath sonicator for 10 minutes before use.

The 500 mg/mL PEG solutions were freshly prepared on the day of sample measurement by adding the suitable molecular weight PEG (8 kDa, 20 kDa, and 35 kDa) to water and gently swirling by hand until a clear solution was obtained. The solution was kept at room temperature until use.

### Synthesis and Preparation of the CrD

The synthesis of CrD (Table 1) was outsourced to Integrated DNA Technologies, Inc., Iowa, using the sequence described by Murade and Shubeita.^31^ The lab-ready CrD stock of 100 µM was diluted per the manufacturer’s recommendation by adding a 1:10 ratio of Tris-EDTA buffer solution. The 10 µM working stocks were stored at −20°C until further use.

During sample preparation, the 10 µM CrD stock was thawed to room temperature and diluted to a final concentration of 0.5 µM CrD in 20 mM HEPES, 150 mM NaCl, pH 7.5, with or without crowders such as Ficoll PM 70 and PEG 8 kDa. Before beginning the measurement, the sample was briefly heated to 90 °C using a dry bath for 10 minutes to disrupt any preformed structures. Then, the sample was allowed to cool to room temperature for at least 20 minutes. Unless otherwise stated, our experiments used 150 mM NaCl to approximate physiological salt conditions.^44^

### Fluorescence Spectroscopy

The fluorescence spectra of 0.5 µM CrH2 and the 0.5 µM CrD sensor were obtained using a JASCO FP-8350 spectrofluorimeter with λ_ex_ = 488 nm, and a 2.5 nm slit width in a 200 µL low-volume quartz cuvette. The emissions were recorded from 500 nm to 700 nm with a 5 nm slit width and a data pitch of 0.5 nm. The integrated area under the curve (AUC) of the donor and the acceptor fluorescence emissions, with a cutoff at 561 nm, was used to calculate the donor/acceptor (D/A) intensity.^45^

### Two-color Fluorescence Microscopy

Either CrH2 or CrD samples were prepared in 20 mM HEPES, 150 mM NaCl, pH 7.5, with 1 mM Trolox and 1 mM DTT, and 0-400 mg/mL PEG or Ficoll. Coverslips (Corning, #CLS2980223) were cleaned in a base-peroxide bath comprised of H_2_O_2_, NH_4_OH, and H_2_O at a 1:1:6 volume ratio at 70 °C for 90 s, then rinsed with water, dried with nitrogen gas (5.0 grade, Airgas), and plasma cleaned in an O_2_ (Industrial grade, Airgas) plasma cleaner (PDC-32G, 115 V, Harrick Plasma) at 140-280 Torr, medium power, for 2 minutes. The coverslips were then used to assemble a sandwiched sample cell of solution between two coverslips with a 13 mm diameter, 0.15 mm depth spacer (Grace Biolabs). Samples were imaged using an Olympus IX-83-based home-built setup. Our microscope has previously been described (Figure 1B) ^46^. Briefly, AcGFP1 and Alexa488 are excited with a 488 nm laser (TOPTICA Photonics). The excitation was reflected by a 488 nm dichroic (ZT488rdc) and focused into the sample in a widefield epifluorescence setting with Olympus UAPON 100X/1.49NA with an approximate power density of 10^4^ 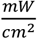. The emission from FRET was collected in inverted epifluorescent mode, split into two detection regions using a 561 nm dichroic (ZT561rdc-UF2), and imaged onto a Photometric Prime 95B sCMOS camera with an integration time of 5-10 ms. The images obtained in TIF format were further processed using Python. The data were averaged over an 800-pixel-by-400-pixel region for both colors at the same detection area.

**Figure 1.**
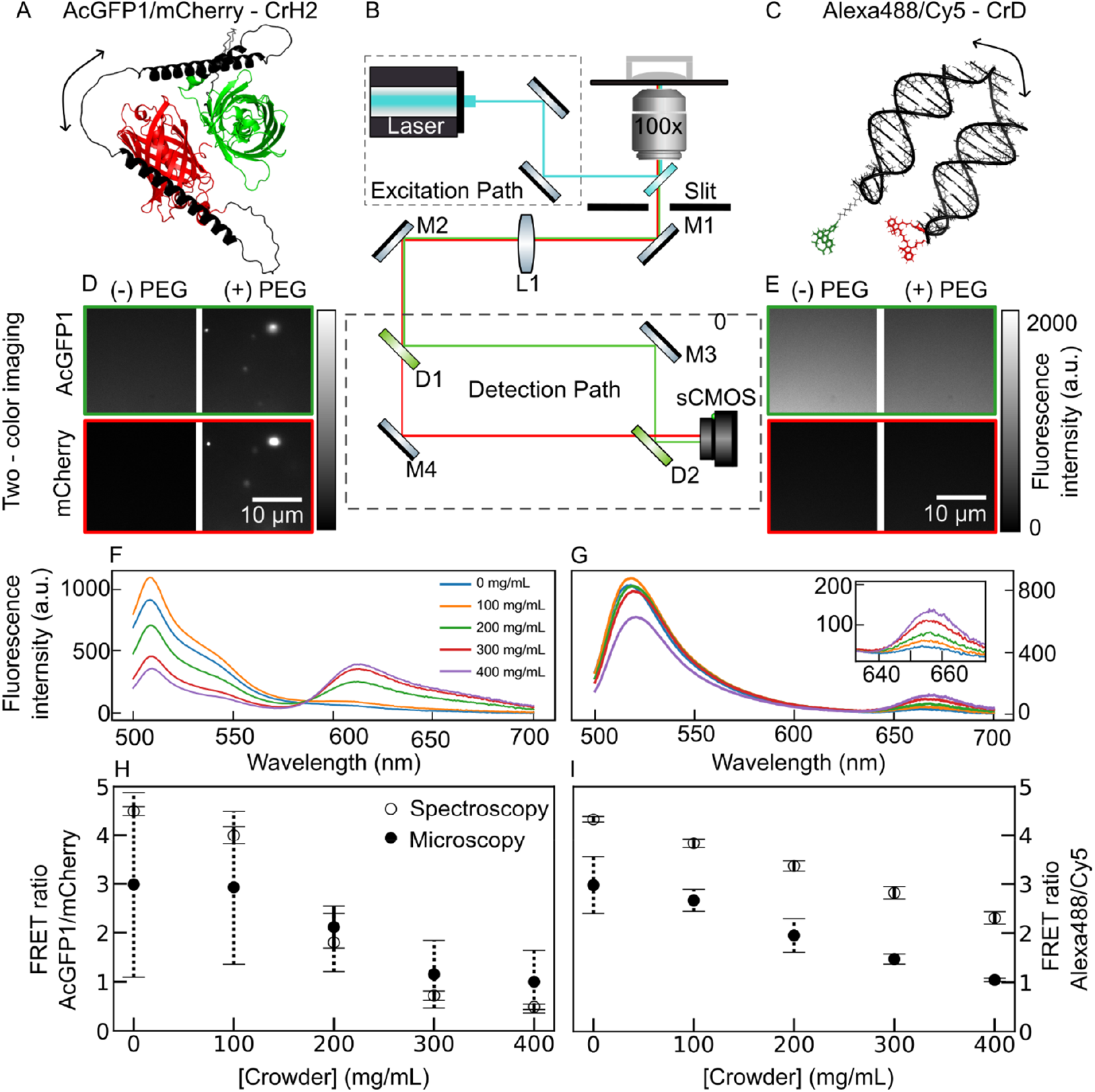
PEG induces a D/A sensor response in CrH2 and CrD, but also microscopic puncta structure formation in CrH2. (A and C) Structure of (A) CrH2 protein (black) with AcGFP1 (green) and mCherry (Red) generated using AlphaFold 3 and (C) Alexa 488(Green)/Cy5(Red)-based CrD, whose backbone was modeled using AlphaFold 3, and later fluorophores were added using UCSF Chimera. Black arrows indicate proposed flexible regions leading to crowding response. (B) Experimental setup of the two-color microscope. See the methods section, “Two-color Fluorescence Microscopy,” for details. The labeled components M1, M2, M3, and M4 are mirrors, D1 and D2 are 561 nm dichroic mirrors, and L1 is the imaging lens. (D-E) The two-color imaging of the (D) 0.5 µM CrH2 sample and (E) the 0.5 µM CrD sensor with and without 400 mg/mL of PEG 8 kDa. (F-G) Fluorescence spectra of the (F) CrH2 and (G) CrD with a 0 - 400 mg/mL titration of PEG 8 kDa. (H-I) The AcGFP1/mCherry and Alexa488/Cy5 (D/A) ratio from two-color microscopy and fluorimeter as a function of 0 – 400 mg/mL of PEG 8 kDa increases for 0.5 µM CrH2 and (I) 0.5 µM CrD. The error is the standard deviation of the mean of at least two measurements.

### Quantification of the CrH2 in the Fluorescence Puncta by Two-color Fluorescence Microscopy

As observed through two-color fluorescence microscopy imaging, we employed intensity-based calculations using MATLAB and Python scripts to obtain relative concentrations of the polydispersed sample spread over a micrometer area, including the droplets. We calibrated our microscope by imaging fluorescence intensity from eGFP and mCherry fluorophores at known concentrations and correlating the intensity to a concentration of the CrH2 fluorophores at a defined laser power. The partitioning coefficient (*K*_*p*_) was calculated by comparing the intensities of the CrH2 samples in the background vs. puncta using the relation *K*_*p*_ = (*I*_*c*_ – *I*_*b*_)/(*I*_*d*_ – *I*_*b*_).^34^ Here, the *I*_*b*_ is the average fluorescence intensity of the blank sample in one frame in the absence of CrH2, *I*_*d*_, and *I*_*c*_ represent the average intensity in one frame of the dilute and condensate puncta structures, respectively. Standard deviations were calculated using at least three different images.

### Fluorescence Recovery After Photobleaching (FRAP)

Samples of 0.5 µM CrH2 in 20 mM HEPES, 150 mM NaCl, pH 7.5, and 400 mg/mL PEG 8 kDa were imaged with a 63x oil-immersion objective (NA 1.4) using a Leica HyVolution SP8 confocal microscope (Leica Microsystems, Germany) equipped with a DMi8 CS motorized stage, a pulsed white light laser, and 2 HyD SP GaAsP and 2 PMT detectors.^34,47–49^ For FRAP, a two-pixel-by-two-pixel region of interest (ROI) was selected within bright puncta areas of the sample, and photobleaching was performed using a 488 nm argon laser with a power of 2.4 mW and an intensity of 80%. The measurements were recorded using LAS X v.3.5 acquisition software for 20 prebleach frames, 100 bleaching frames, and 600 postbleaching frames (1.3 s/frame). The recovery intensity is normalized with the intensity before bleaching and then is analyzed using Origin software. The recovery data was analyzed with the exponential decay equation,^34^ I_normalized_ A(1 − *e*^−*bt*^) + *C*.

### Far-UV Circular Dichroism (CD)

CD spectra were acquired on an Applied Photophysics Chirascan Circular Dichroism Spectrometer in 20 mM sodium phosphate buffer, pH 7.0, using a 0.1 mm quartz cuvette. The data was acquired in the absence and presence of PEG 8 kDa with either 0 or 150 mM NaCl using a 1 nm data pitch at 25℃ from 190 – 280 nm. The represented spectrum is an average of two measurements acquired on different days. The secondary structure deconvolution was performed using the online server Dichroweb and CONTIN analysis program and SP175t dataset (data not shown).^50,51^

## Results

### Comparison of FRET Response from CrH2 vs CrD

The design philosophies of protein- and DNA-based molecular crowding sensors dictate their application, efficiency, and robustness. First, CrH2 is based on the sensor originally proposed by Boersma *et al*.^26^ and later modified by Sukenik *et al*..^12,29,32^ The two fluorophore domains in CrH2 are separated by a linker domain containing a flexible glycine-serine-glycine (GSG)_6_ repeat unit along with a rigid alpha-helical stretch based on glutamic (E) and lysine (K) residue repeats spaced by alanine, where helicity is anchored by ion pairs between anionic E and cationic K at pH 7.5.^52^ The energy transfer between the AcGFP1/mCherry leads to a proposed continuous change in donor to acceptor ratios, D/A, to act as a molecular conformational “rheostat” for crowding.^29^ Similar to the proposed design, the AlphaFold prediction of the CrH2 protein structure renders two structurally distinct AcGFP1 and mCherry domains separated through a linker polypeptide region with two helical motifs (Figure 1A).^53^ We also note that FRET-based sensors are often purified with an N/C-terminal His_6_ tag, and studies oftentimes do not remove the purification tag.^20,26,27^ The contribution of the known self-assembly-promoting poly-His region^54^ is unclear, but the given the short sequence and large distance from the crowding-responsive helices, it is assumed not to interfere with crowding response.

Next, we modified a DNA-crowding sensor described by Murade *et al*. to accommodate the spectral range of our microscope (Table 1, Figure 1C and S4).^31^ The CrD comprises a 59-base pair single strand of DNA, proposed to form two double-stranded arms, with the donor and acceptor organic fluorophores at the 14th and 46th base pairs, respectively. The original probe, labeled with Cyanine3 (Cy3)/Cyanine5 (Cy5) would render crosstalk-based responses on our home-built microscope, which features a 488 nm excitation laser and a 561 nm dichroic to split emission for two-color microscopy imaging (Figure 1B, S4).^31^ To broaden the usability of the DNA-based crowding sensor, we incorporated Alexa 488/Cy5 as a FRET pair, while maintaining the same oligonucleotide backbone sequence (Figure 1C).^31^ The Alexa 488 donor fluorophore has a higher quantum yield than Cy3. Still, it has less spectral overlap with the Cy5 acceptor pair, resulting in comparable responses to crowding between our Alexa 488/Cy5 sensor and the original Cy3/Cy5 (Figure S5). Thus, we have modified and developed CrD, a blue laser-compatible (488 nm) Alexa 488/Cy5 DNA-based crowding sensor. Therefore, our sensor can be easily implemented on microscopes designed with optics for common fluorescent proteins mCherry and GFP. A comparison of the spectral properties of our sensor with those reported by Murade *et al*. is presented in Figure S4 and Table S2.

Fluorescence spectroscopy of CrH2 shows a FRET response to PEG-induced crowding (Figure 1F). We observe a decrease in the ratio of donor to acceptor fluorescent intensity (D/A) with increasing concentrations of PEG 8 kDa (0 – 400 mg/mL) (Figure 1F). A higher D/A indicates that donor and acceptor are held apart; this is attributed to a more conformationally expanded state of the protein. A decrease in D/A with the addition of crowder indicates that the donor and acceptor are closer, leading to an assumed intramolecular FRET response to increased crowding. Generally, a linear decrease (Figure S6) in the D/A fluorescence intensity ratio is expected with the addition of crowders.^26^ Yet, we observe non-linear response of CrH2 with a more drastic change in D/A between 100 mg/mL and 200 mg/mL that we investigated further via ensemble spectra and microscopy.

PEGs are known to interact with proteins^35^ and alter their fluorescence response.^55,56^ More recently, PEG has been shown to influence the fluorescence lifetime of eGFP molecules.^55^ Inspecting CrH2 emission in the presence of PEG 8 kDa (Figure 1F), we observed an unexpected initial increase in donor intensity at 100 mg/mL PEG, followed by a subsequent decrease in donor intensity at concentrations greater than 100 mg/mL PEG, compared to the protein alone sample. In an independent titration experiment with PEG 8 kDa and Ficoll PM 70, we see an increase in the fluorescence emission intensity of a commercially available eGFP molecule (Figure S7A, B). The eGFP is a close approximation of AcGFP1 due to its close sequence homology and fluorescence properties.^57,58^ The photophysical mechanisms underlying the elevation of fluorescence intensity of eGFP in the presence of PEG are not well understood. However, it has been argued to stem from the change in the refractive index in the close vicinity of the fluorophore.^55,59^ In addition, we did not observe appreciable fluorescence sensitivity of mCherry in the presence of PEG 8 kDa and Ficoll PM 70 (Figure S7C, D), consistent with previous reports.^56^

The average CrH2 sensor response to crowding from PEG is consistent between ensemble fluorometry and the average intensity within the nanoscale fluorescence microscopy (Figure 1H). The D/A values obtained from the fluorimeter closely correlate with the average D/A intensity from microscopy imaging taken over an 8 µm x 5 µm area, with a decrease in the D/A trend in the FRET response to the crowding environment induced with PEG. The micrograph from the two-color microscopy imaging of CrH2 is shown in Figure 1D, which was acquired in the presence of 400 mg/mL of PEG 8 kDa and shows an average gain in intensity in the acceptor channel when the entire image is considered, as reflected in the D/A profiles (Figure 1H).

Yet, as we titrate CrH2 samples with PEG, we observe distinct, fluorescently active, round puncta in both donor and acceptor channels (Figure 1D) and a subsequent decrease in the fluorescence surrounding the puncta (Figure 1D). We also observed the fluorescent puncta at low salt conditions of 20 mM sodium phosphate, pH 7.5 buffer recipe (Figure S8). No such puncta are observed in the CrD samples (Figure 1E), which further shows a linear response to PEG-induced crowding (Figure 1G). Additionally, no puncta are observed with the CrH2 protein alone (Figure 1D, (-) PEG condition) or fluorescent protein donor or acceptor alone controls (Figure S9). In the next section, we systematically investigate the increased CrH2 fluorescence in the puncta region vs the surroundings and the transition thereof, leading to phase separation and puncta formation.

### CrH2 Fluorescent Puncta Formation in the Presence of Polyethylene Glycol is Concentration-Dependent

High-resolution nanoscale images of the CrH2 reveal spatial details that are missed during ensemble fluorometry measurements of *in vitro* crowding (Figure 2). We observe fluorescent puncta in the two-color microscopy images of 0.5 µM CrH2 samples at or above 200 mg/mL of PEG 8 kDa (Figure 2B).^26^ The fluorescent puncta were not seen in the absence of crowder (0 mg/mL, Figure 2A, B) or in the presence of Ficoll PM 70 (0 – 400 mg/mL, Figure 2A). The puncta can be seen on donor and acceptor channels, suggesting enrichment of CrH2 from the surrounding and into the puncta. We also rarely observe fluorescence puncta at 100 mg/mL of PEG 8 kDa (data not shown), suggesting microinhomogeneities could occur, which may not accurately represent the entire sample. Interestingly, we did not see any puncta with Ficoll PM 70 as a synthetic crowder (Figure 2A), suggesting this is solely PEG-induced behavior.^13,60^

**Figure 2.**
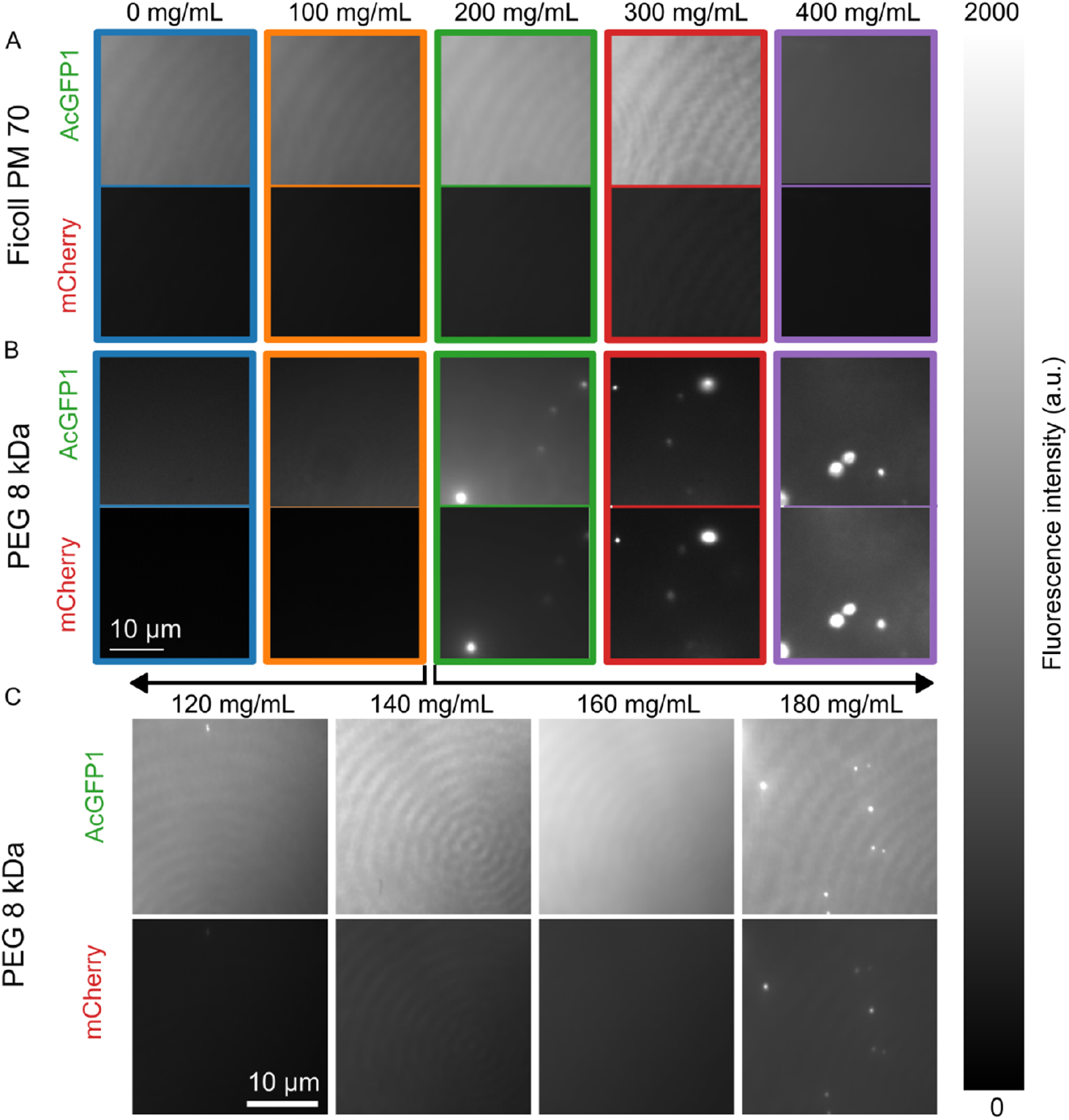
Ficoll PM 70 does not induce fluorescent puncta formation of CrH2, whereas PEG 8 kDa does. Two-color imaging of the 0.5 µM CrH2 with Ficoll PM 70 or PEG 8 kDa at 0 – 400 mg/mL crowder concentrations in the buffer 20 mM HEPES, 150 mM NaCl, pH 7.5, 1 mM DTT, and 1 mM Trolox. (A) Micrographs show Ficoll PM 70 does not induce fluorescent puncta formation of CrH2 at 0 – 400 mg/mL Ficoll PM 70 concentrations. (B) Micrographs show the CrH2 phase separates at or above 200 mg/mL concentrations of PEG 8 kDa. (C) Investigation into the critical PEG 8 kDa concentration required to induce fluorescent puncta formation at 0.5 µM CrH2. Probing at 20 mg/mL steps of PEG 8 kDa, we see a prominent increase in the incidence of fluorescent puncta at 180 mg/mL of PEG 8 kDa. We performed at least two independent experiments with at least three random fields of view captured at each concentration of PEG and Ficoll. A representative micrograph is presented for each concentration of the crowder.

Next, to identify the minimum critical concentration of PEG 8 kDa required for puncta formation, we titrated 0.5 µM CrH2 with increasing PEG concentrations in finer steps of 20 mg/mL from 120 to 180 mg/mL (Figure 2C). The crowder titration was performed, while keeping the buffer components identical, except for the crowder as the titrant. As we scan through the PEG concentrations and reach the 180 mg/mL threshold, we see a substantial increase in puncta observed in the two-color images (Figure 2B). After this crowder concentration is reached, the size of the fluorescent puncta increases further as a function of PEG concentration (Figure 2B). We also observe the early occurrence of fluorescent puncta, even at a 140 mg/mL concentration of PEG 8 kDa, when a higher concentration of 5 µM CrH2 was used (data not shown). Thus, the fluorescence puncta are formed as a function of both PEG 8 kDa and CrH2 concentrations.

### CrH2 has Liquid-like Properties Inside the Fluorescent Puncta

The observed CrH2 puncta are in a liquid state as characterized by fluorescence recovery after photobleaching (FRAP). We performed ultracentrifugation on a freshly thawed CrH2 aliquot to eliminate the possibility of preformed aggregates in our samples. We used the supernatant for sample preparation of a 5 µM CrH2 sample in the presence of 400 mg/mL PEG 8 kDa. We observe fluorescence recovery over time in the photobleached area (Figure 3B), indicating the transfer of fluorescently active molecules into the bleached region, which confirms that the punctate structures we observe are liquid condensates. We quantify the recovery in a 2-pixel by 2-pixel region from three different puncta and fit the recovery data points with an exponential model.

**Figure 3.**
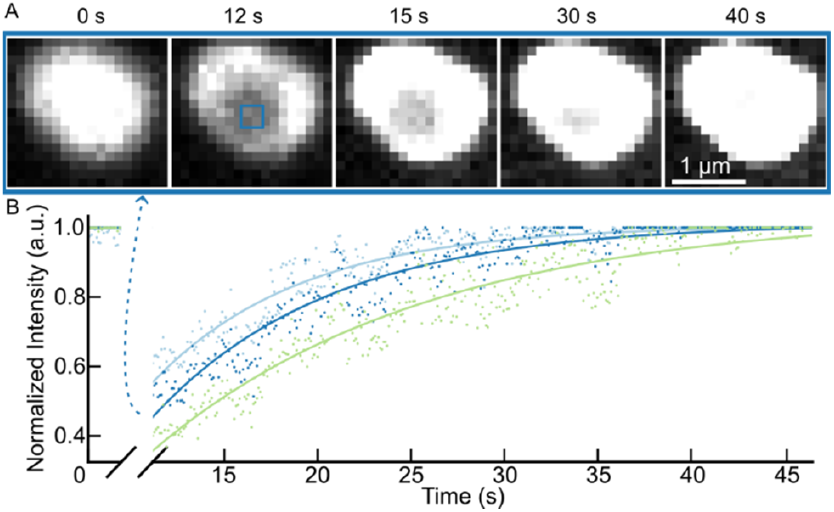
The CrH2 condensates have liquid-like FRAP recovery profiles. (A) The prebleach (0 s), bleach (12 s), and postbleach (after 15 s) of one representative condensate from a 5 µM CrH2 sample in the presence of 400 mg/ml PEG 8 kDa are shown. The data points are averaged from a 2-pixel by 2-pixel region (blue square box at 12 s). (B) The FRAP data points are fitted with an exponential model (solid line). The variation present in recovery for three different puncta is shown in B. The light blue, dark blue, and green scatter points, along with the fit lines, indicate data and fit lines from three puncta, respectively, with the dark blue data shown in detail in (A) as indicated by the dashed arrow.

### PEG Induces Enrichment of CrH2 Inside Fluorescent Puncta

Spatial information in two-color microscopy allows us to understand the PEG-induced phase transition of CrH2 quantitatively. We measured the relative change in fluorescence intensity within the puncta structures formed upon the addition of 0 – 400 mg/mL PEG 8 kDa with 0.5 µM CrH2. We observe an increasing trend in the acceptor channel fluorescence intensity from puncta formed with 0 – 400 mg/mL of PEG 8 kDa (Figure 4). The increase in the acceptor fluorescence signal may be attributed to two competing processes: 1) accumulation of the CrH2 molecules into the puncta, and 2) crowding-induced FRET-based enhancement of acceptor fluorescence. However, the donor channel fluorescence intensity shows a steadier decrease, followed by enhancement only at the highest concentration of the crowder (Figure 4A). The total donor channel fluorescence arises from opposing processes. One, from FRET-based quenching of donor fluorescence, in parallel with the fluorescence enhancement due to the accumulation of CrH2 molecules into the puncta, we see a steady net decline in intensity from 0 – 300 mg/mL of PEG 8 kDa. However, at the highest 400 mg/mL concentration of PEG, we observe the highest increase of donor fluorescence, suggesting a depletion of CrH2 in the dilute phase and sequestration of the sensor into the puncta (Figure 4A).

**Figure 4.**
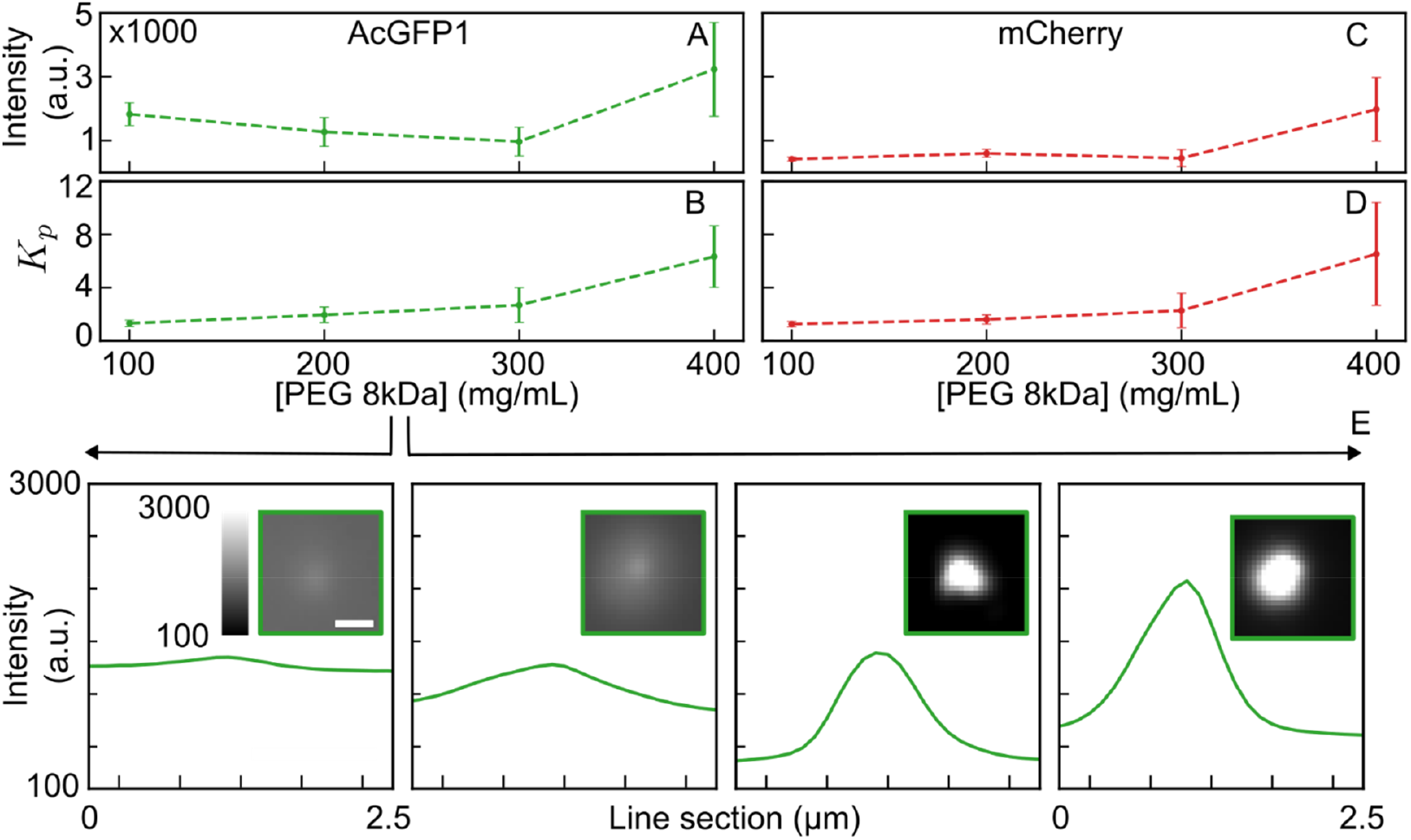
PEG 8 kDa induces CrH2 enrichment into the fluorescent puncta. Data from two-color imaging of the 0.5 µM CrH2 with PEG 8 kDa at 0 – 400 mg/mL concentrations in the buffer 20 mM HEPES, 150 mM NaCl, pH 7.5, 1 mM DTT, and 1 mM Trolox. (A) Donor and (C) acceptor channel fluorescence intensity inside the puncta as a function of 0 – 400 mg/mL PEG concentrations. (B) donor channel and (D) acceptor channel specific partition coefficient (K_*p*_) across dilute and dense phases, as a function of 0 – 400 mg/mL PEG 8 kDa concentrations. (E) The intensity line profile originates from the donor channel micrograph of a 0.5 µM CrH2 sample in the presence of 100 - 400 mg/ml PEG 8 kDa. It represents an average intensity from 25 vertical line-sections of the boxed inset with a representative enlarged fluorescence puncta. The inset shows a 25-pixel by 25-pixel area of the sample, with an 800 nm scale bar.

The partition coefficient quantifies the increase in CrH2 within the puncta as crowder concentration increases. The partition coefficient (*K*_*p*_) was calculated by measuring the fluorescence intensity from the background dilute phase (*I*_*b*_) and fluorescent puncta (*I*_*c*_) to determine the relative distribution of CrH2 molecules using methods described elsewhere (Figure 4E).^34^ Interestingly, we see an increase in the *K*_*p*_ across the 0 – 400 mg/mL PEG 8 kDa samples in both donor and acceptor channels (4B and 4D). The statistically identical increase of *K*_*p*_, which is a relative measure of partition coefficient, for both the donor depletion in the dilute phase.

### The Excluded Volume Effect is Not the Singular Driving Force Behind CrH2 Phase Separation

The crowder type, size, and excluded volume impact the extent of CrH2’s FRET response (Figure 5A). We have compared the excluded volume effect of Ficoll PM 70 and PEGs ranging from 1–35 kDa as crowders on CrH2. Using our two-color microscope setup (Figure 5A), we measured the ratiometric changes in AcGFP1 vs. mCherry channel emission intensities of CrH2 in both crowder types as a function of crowder concentration (Figure 5B). PEGs of different sizes confer a phase transition of CrH2 into fluorescent puncta. Two-color imaging (Figure 5A) shows that PEGs of 1, 20, and 35 kDa can also induce phase separation of 0.5 µM CrH2 into condensates, similar to the PEG 8 kDa. When compared to PEGs of various sizes at the same concentration, Ficoll PM 70 exhibits a moderate response, with a 1.2-fold reduction in the D/A ratio at 400 mg/mL. We have taken the hydrodynamic radius of each polymer from the literature and, approximating the size as are hard spheres, calculated the volume of each particle 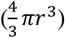 and subsequently the excluded volume fraction using empirical volume calculations (Table S4).

**Figure 5.**
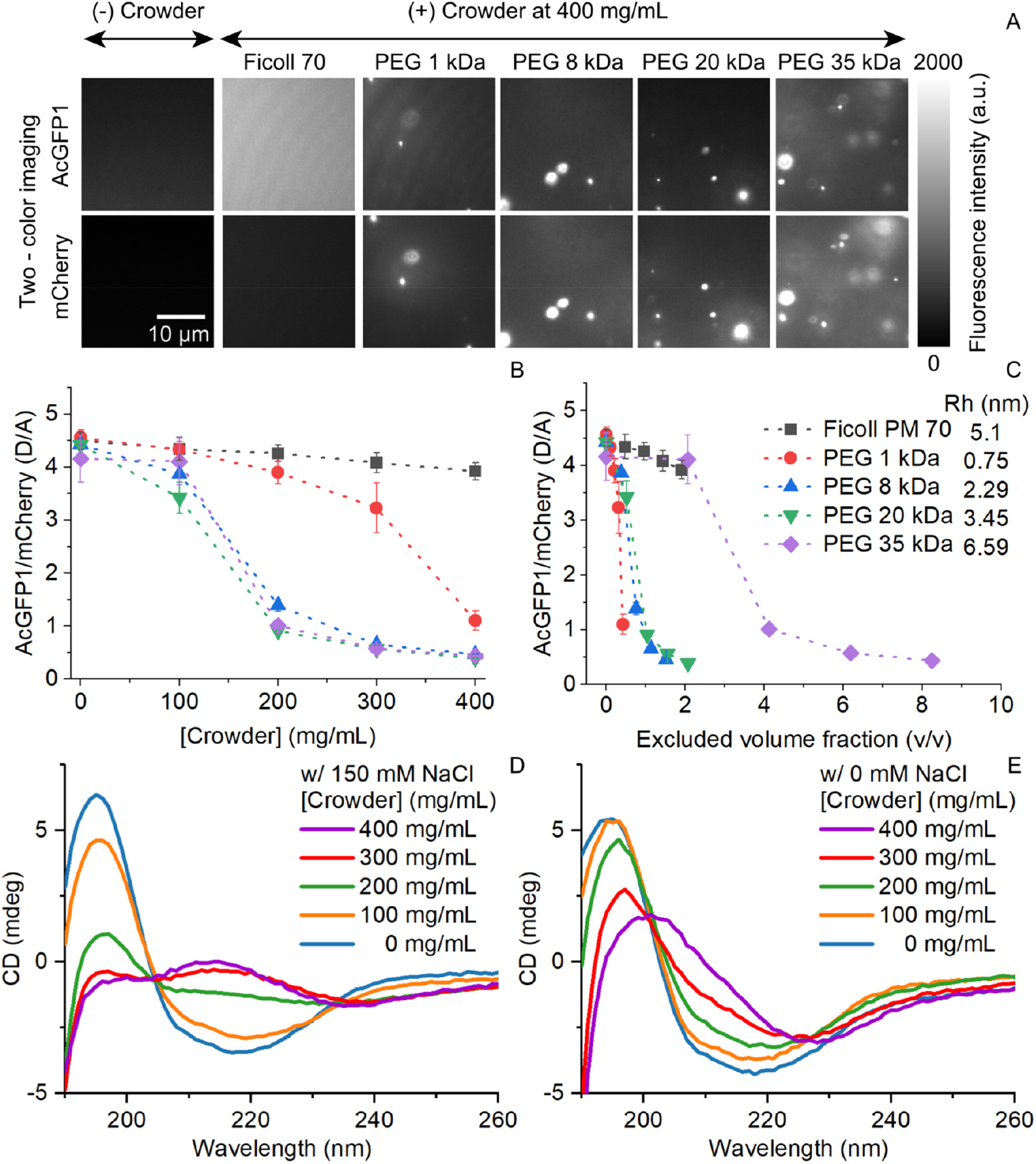
The excluded volume effect is not the singular driving force behind CrH2’s FRET response. (A) PEGs of 1-35 kDa sizes induce phase separation of 0.5 µM CrH2 at higher crowder concentrations. The two-color imaging of a µM CrH2 sample, without or with 400 mg/mL of Ficoll PM 70 and PEG of different sizes. (B – C) The fluorometry AcGFP1/mCherry (D/A) ratio of 0.5 µM CrH2 sample with Ficoll PM 70 and PEGs 1 – 35 kDa at 0 – 400 mg/mL concentration vs (A) [crowder] (mg/mL) and (B) excluded volume fraction (v/v). The error is the standard deviation of the mean of two measurements. The dashed lines connect the symbols for illustrative purposes only. The hydrodynamic radius (Rh) values are from Table S4. (D-E) CD spectra of 4 µM CrH2 in 0 – 400 mg/mL PEG 8 kDa (D) with 150 mM NaCl and (E) with 0 mM NaCl.

CrH2 has a non-linear FRET response to PEGs of various sizes. The fluorometry FRET response in the presence of Ficoll PM 70 with a 5.1 nm hydrodynamic radius is linearly correlated across 0 – 400 mg/mL concentration of crowder with D/A ratio ranging from 4.5 ± 0.2 to 3.9 ± 0.2, respectively (Figure 5B). The PEG 1 kDa has a smaller hydrodynamic radius of 0.75 nm; yet, it has a larger impact on the D/A, which decreases 4.2 times from 4.6 ± 0.1 to 1.1 ± 0.2 across 0 – 400 mg/mL of crowder (Figure 5B). With larger PEGs of 8, 20, and 35 kDa, having hydrodynamic radii of 2.29, 3.45, and 6.59 nm, respectively, the D/A decreases by 9.6, 11.3, and 9.5 times, respectively, from concentrations of 0 mg/mL to 400 mg/mL of the crowders (Figure 5B). The D/A ratio of the PEGs 8, 20, and 35 kDa reduced from 4.4 ± 0.1 to 0.4, 0.4, and 0.4, respectively.

Next, to understand whether the excluded volume conferred by the Ficoll and PEGs is the primary driving force for CrH2’s FRET response, we calculated the excluded volume fraction occupied by the crowders. The excluded volume is the ratio of the volume occupied by the total amount of crowder molecules (V_ev_) in a solution to that of the volume of solution (V_t_) (Figure 5C, Tables S3 and S4). The smaller-sized PEG 1 kDa, with the lowest excluded volume fraction, has the largest number of ethylene glycol (EG) subunits in the solution at a crowder concentration (mg/mL) (Tables S3 and S4), and can confer a comparable fold change to the D/A, as that of the larger PEG size. The larger PEGs confer a comparable fold change in the D/A response, although the excluded volume fraction increases as a function of PEG size (Figure 5B, C). However, Ficoll PM 70, with a larger average molecular weight (70 kDa) and a 5.1 nm hydrodynamic radius, is unable to confer a comparable D/A change compared to PEGs of similar hydrodynamic radius. Thus, the excluded volume effect alone does not account for CrH2’s FRET response with PEG.

PEG may decrease the alpha helical content of CrH2 to drive phase separation. We use circular dichroism (CD) spectroscopy to assess the secondary structure of CrH2, which contains both beta-sheet containing fluorescent proteins^61^ and the alpha helical hinge region(Figure 5 D, E).^12^ In the absence of PEG, the CrH2 spectra have signatures of both the alpha and beta portions, with a positive peak at 195 nm and a broad negative region spanning 210-230 nm.^62^ Quantitative deconvolution algorithms such as BestSel^63^ and Dichroweb^51^ result in secondary structure content that does not align with the protein design and disagree with one another, likely due to the challenging overlap of the mixed structural content and the changing solvent conditions upon condensation. Therefore, we focus on qualitative interpretations of the spectra. In the absence of PEG, the CrH2 spectra in buffer with 150 mM NaCl have signatures of both the alpha and beta portions, with a positive peak <200 nm and a broad negative region spanning 210-230 nm. As 8 kDa PEG is added to CrH2, a drastic change in the spectra occurs between 100 mg/mL and 200 mg/mL, agreeing with the phase transition point in microscopy. At ≥200 mg/mL 8 kDa PEG, the negative baseline and loss of intensity in the near-UV indicate that the phase separation-induced turbidity at higher crowder concentration limits the acquisition of better CD spectra. Therefore, we obtained CD spectra of CrH2 and 8 kDa PEG in the absence of NaCl (Figure 5E), as it is known that achiral buffer components, such as salt (Cl^-^), absorb light, reducing the photons available for the protein to absorb.^63,64^ The spectra in 0 mM NaCl show more interpretable spectral signatures even at 400 mg/mL 8 kDa PEG. Notably, we observe a loss of the 208 nm peak, indicating a decrease in alpha helical content. At PEG concentrations >200 mg/ml, we observe a red shift of the minimum at 220 nm. This shift is often observed in membrane proteins with beta sheet structure embedded in a detergent micelle or a phospholipid bilayer and arises due to the change in dielectric constant in the hydrophobic core compared to water.^65^ Therefore, we interpret our spectra as arising from beta proteins buried in the hydrophobic core of the condensate. At 400 mg/mL, this peak is due to the remaining beta barrels of the fluorescent proteins in CrH2 that we observe in microscopy.

## Discussion

A lack of understanding of the behavior of protein crowding sensors under phase-separation conditions prompted us to explore the material conditions that drive the formation of liquid phase CrH2 and the corresponding sensor response. Synthetic crowders, such as PEG, are believed to impart excluded volume effects, enabling a crowding-induced compaction-driven FRET response from the crowding sensor CrH2. PEG has been used as a synthetic crowder with crowding helix-based protein sensors^26,32^ and with other biomolecular crowding sensors.^66^ Yet, PEG has also been used to drive phase separation of proteins for separation purposes and is known to interact with protein surfaces. We observe that the PEG-induced phase separation is not solely driven by excluded volume; chemical effects of the crowder must also be considered.

Fluorescence microscopy sheds light on the widely underexplored behavior of the CrH2 crowding sensor, examining the nature of the condensates at the microscale. The bulk response observed in Figure 1H could easily be used as a calibration with a linear regression fit, providing a similar R^2^ from both fluorimeter and microscopic measurements (Figure S6). Without knowledge of the phase separation occurring, the response could falsely be used to report on excluded volume effects of an assumed single-phase system. Due to the transient and fragile nature, along with diverse size ranges of condensates, it is challenging to characterize them. Therefore, fluorescence microscopic imaging techniques are employed to understand the droplets.^67^ Only the nano- to micrometer-scale spatial information provided by two-color fluorescence microscopy (Figure 2) and FRAP (Figure 3) reveals the presence of liquid-phase-separated droplets. Further microscale spatial analysis within and around the droplet reveals spatial alterations in concentration across the dilute phase and enrichment of the probe in the condensed phase (Figure 4). Therefore, the average sensor response reports on two distinct phases and hence, two different levels of crowding.

The FRET of a biomolecular sensor undergoing condensation can complicate the signal readout. While FRET-based protein crowding sensors have been shown to form coacervates,^26^ they have still been used in studies to understand the physicochemical properties of membraneless organelles.^27^ In an earlier report, a well-established fused in sarcoma (FUS) protein, an ALS marker that undergoes phase separation, has been shown to drive a C-terminal eGFP carrier tag into the condensate.^68^ Furthermore, the eGFP tag in the condensed phase underwent homoFRET. The intermolecular energy transfer between two chemically and spectrally identical eGFP molecules is facilitated due to the inherent minimal Stokes shift in the excitation and emission spectra. Currently, anisotropy is the only available biophysical tool to differentiate between intermolecular homoFRET, intermolecular hetero-FRET, or intramolecular hetero-FRET^69^ that could occur with CrH2 in the dense, condensed liquid phase. Future work to understand the CrH2 sensor FRET response within condensates, via homo- or hetero-FRET, could pursue these approaches. Time-resolved measurements^70^ could further reveal the dynamics of the individual AcGFP1 and mCherry fluorescent proteins in the presence of PEG that led to our observed increased emission of GFP (Figures 1F, S7), and are convolved with the sensor response.

The role of the chemical makeup and structure of both proteins and PEG in phase separation has been explored. The mechanistic understanding of PEG-driven phase separation of proteins has identified a broad range of driving forces, including electrostatic, hydrophobic, and hydrogen bonding, or a combination of these, leading to phase separation, which is further protein-dependent.^35,71,72^ Condensation in the binary mixture of CrH2 and PEG in a buffered solution could be explained by interactions between the solvent-exposed helical regions and an increase in the effective protein concentration as the driving force of phase separation.^34^

First, CrH2 being a chimeric protein, the β-sheet rich fluorophore domains are connected through a flexible linker region enriched in (GSG)_5_-rich disordered and (EAAAK)_6_-rich helical regions.^29^ We observe CrH2 fluorescence at all concentrations of PEG, and our control spectroscopic measurements (Figures S7, S9) demonstrate that PEG does not significantly influence the β-sheet domains of GFP or mCherry alone. The flexible linker, rich in α helix, is designed to remain solvent-exposed, allowing the fluorophore domains to remain apart in solution. However, our CD spectra of CrH2 indicate structural changes in the presence of 8 kDa PEG (Figure 5D, E). In earlier work^29^ and in the current study (Figure S3B), a CD-melt analysis of CrH2 reveals a slight decrease in the α-helical content (CD at 222 nm) over elevated temperatures, suggesting a weakening of the helix.^29^ Recent advancements have revealed the role of salt in destabilizing helical peptides.^73,74^ Interestingly, the crystal structure of another fusion protein with a short helical-linker motif shows a broken helix in the presence of 20 % (w/v) PEG 6 kDa and 0.2 M CaCl_2_.^75^ In addition, short 2x or 3x repeats of (i+3)_EK_ motifs have been found to lack α-helical structure.^76,77^ Linker domains in fusion proteins with a solvent-exposed helical motif stabilized by salt bridges are prone to charge screening. The melted short-helical linkers can participate in intermolecular interactions, thereby favoring multimers.^76^ Therefore, a thorough understanding of “rigid” and “stable” helices in the presence of physiological salt levels, crowders, and temperatures should be considered in sensor design.^52,78^

Second, PEG is a known inducer of phase separation through facilitating intermolecular interactions.^35,37^ PEG also increases the effective concentrations of proteins,^79^ and enables intermolecular noncovalent interactions that lead to self-assembly. Additionally, proteins with helical linkers may also encounter a PEG-induced dielectric decrement due to dehydration.^80^ The sequence-specific molecular grammar in proteins makes weak, multivalent, intermolecular, and protein-solvent interactions necessary for phase separation.^35,81,82^ The saturating protein concentration, pH, and salt conditions can promote inter-protein interactions and decrease the entropy (ΔS) necessary for phase separation.^35,83^ A mechanistic understanding of phase separation induced by PEG is shown to outweigh the excluded volume effects as the only driver (Figure 5).^34^ Moreover, PEG is shown to co-condense with proteins and induce homotypic, heterotypic, and segregative phase separation effects.^34,84,85^

Although it is difficult to predict what specifically drives the phase separation behavior of CrH2 (refer SI section “The Sequence Properties Potentially Drive the Phase Separation of CrH2” and Figure S10), chemical interactions between the flexible helical region and PEG, as described above, may be the reason behind our observations. We believe that the helix-spaced-disordered framework in CrH2, in the presence of PEG and physiological salt concentration, overcomes the energetic barrier to participate in phase separation via both enthalpic chemical interactions and entropic steric effects.

Overall, selecting the correct type of crowding sensor and soluble crowder for the physical phenomena of interest – either excluded volume macromolecular crowding or intermolecular forces driving phase separation – and performing appropriate control measurements are critical to obtaining a correct analytical readout. We observed a single phase and linear response with the CrD using Ficoll and PEG (Figure 1G, in contrast to CrH2 with PEG), which may be more appropriate for *in vitro* crowding studies with PEG. Yet, single-stranded DNA has also been observed to undergo phase separation in different poly-l-lysine crowders,^13,86^ again pointing to the need for a comprehensive understanding of crowder response in heterogeneous environments. CrH2 could also still be used before its phase transition, as we observed a linear response in lower concentrations of PEG, consistent with previous observations.^26^

We further recommend that microscopy should be used to confirm that phase separation is not occurring for more complex studies, such as those using crowder cocktails with a mixture of sizes to mimic the complexity of cellular environments,^87,88^ or with cellular imaging. Beyond the *in vitro* work which we focus on here, the CrH2 motif was developed to be expressible in living cells.^26^ Little control of sensor concentration levels may be possible through transient transfection,^89^ so subcellular microscopy with punctilious review of the images should confirm that no condensates are observed. Our data also suggest that cellular chemical interactions that do not originate from condensation might impact intracellular crowder readings. Thus, users of this and other biological sensors must be careful to consider the impact of chemical interactions on their readouts.

## Conclusions

Our results provide a guide for selecting and calibrating fluorophore and biomolecular sensors for *in vitro* crowding studies using fluorescence microscopy. We tested FRET-based biomolecular crowding sensors with either a fluorescent protein-alpha helical hinge or synthetic fluorophores and DNA design. Using two-color microscopic imaging, FRAP, and spectroscopic measurements, we demonstrate that CrH2 undergoes *in vitro* phase separation in the presence of PEG and exhibits an anomalous fluorescence sensor response. Our data clarify the differences in crowding responses between CrH2 using PEG vs Ficoll. We have investigated the nature of the fluorescent puncta formed by the former molecular crowder, which we did not observe with the latter. We have used FRAP to decipher the liquid-like properties of CrH2 sequestered inside the fluorescent puncta. A partition coefficient analysis quantified the phase transition of CrH2 from a dilute to a dense, fluorescent puncta. A comparison of excluded volume effects conferred by Ficoll and various size ranges of PEGs enables us to understand that non-singular driving forces influence the phase regime of the CrH2 FRET response. We hypothesize that the phase transition is driven by a combination of entropic volume exclusion and enthalpic effects of the PEG interacting with the alpha-helical domain of the CrH2, which could be a future focus for polarized and time-resolved fluorescence imaging.

## Data, Materials, and Software Availability

All data is available within the article and supplementary materials. Python codes for processing the images may be shared upon request.

## Author Information

Notes: The authors declare no competing financial interest.

## Acknowledgments

We acknowledge funding from the National Institutes of Health Grant R35GM142466 and the National Science Foundation CAREER award MCB-2338323. The authors acknowledge Dr. Rajesh Ramachandran for our research collaborations and for facilitating the use of the Department of Physiology and Biophysics’s ultracentrifuge in the current study. We thank Dr. Andy Liu and Dr. Michael Jenkins for helping us train and use the Leica SP8 confocal microscope at the SOM Light Microscopy Core Facility. The NIH-shared instrumentation grant, S10-OD024996, supported the acquisition of the Leica SP8 confocal microscope. This research made use of the Applied Photophysics Chirascan in the Chemical and Biophysical Instrumentation Center at Yale University (RRID:SCR_021738).We thank Dr. Divita Mathur and Akeshi Aththanayake for assistance with Figure 1C and the Kisley research group for helpful discussion. MAB was partially supported by a National Science Foundation Graduate Research Fellowship under grant DGE-2139841.

## Author contributions

LK, AM, and JA designed research; JA, MAB, and AM performed research; LK, CMD, AM, JA, and DRL analyzed data; AM, JA, and LK wrote the manuscript. All authors edited and revised the paper.

## TOC FIGURE

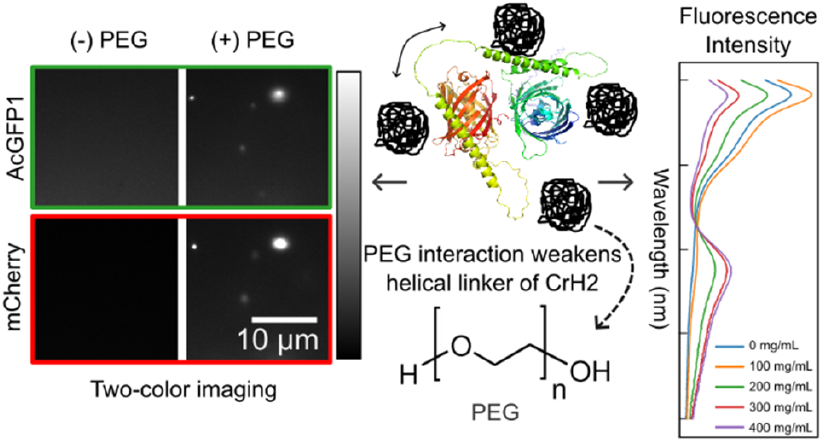

