## Supplemental Text and Figures for "Chemical interactions in polyethylene glycol-induced condensates lead to an anomalous FRET response from a flexible linker-fluorescent protein crowding sensor"

**Amino-acid and nucleic-acid sequences of the CrH2 construct**

The CrH2 probe design was adopted from elsewhere[1,2] and the color-coded amino acid sequence is depicted below. The DNA sequence of the vector containing CrH2 follows the protein sequence.

**Protein Sequence**

MDYKDDDDKGSGSSHHHHHHSSGLVPRGSMVSKGAELFTGIVPILIELNGDVNGHKFSVSGEGE
GDATYGKLTLKFICTTGKLPVPWPTLVTTLSYGVQCFSRYPDHMKQHDFFKSAMPEGYIQERTIFF
EDDGNYKSRAEVKFEGDTLVNRIELTGTDFKEDGNILGNKMEYNYNAHNVYIMTDKAKNGIKV
NFKIRHNIEDGSVQLADHYQQNTPIGDGPVLLPDNHYLSTQSALSKDPNEKRDHMIYFGFVTAAA
ITHGMDELYKTLGMDELYKGSGGSGGSGGSGGSGGSGAEAAAKEAAAKEAAAKEAAAKEAAA
KEAAAKAGSGGSGGSGGSGGSGGSGAEAAAKEAAAKEAAAKEAAAKEAAAKEAAAKAGSGG
SGGSGGSGGSGGSGVSKGEMVSKGEEDNMAIIKEFMRFKVHMEGSVNGHEFEIEGEGEGRPYEG
TQTAKLKVTKGGPLPFAWDILSPQFMYGSKAYVKHPADIPDYLKLSFPEGFKWERVMNFEDGGV
VTVTQDSSLQDGEFIYKVKLRGTNFPSDGPVMQKKTMGWEASSERMYPEDGALKGEIKQRLKL
KDGGHYDAEVKTTYKAKKPVQLPGAYNVNIKLDITSHNEDYTIVEQYERAEGRHSTGGMDELY
K

Number of amino acids: 636

Molecular weight: 67847.41 (~68 kDa)

**DNA Sequence**

atggattacaaggatgacgacgataagggatccggcagcagccaccaccaccaccaccacagcagcggtctggtgccacgcggtagcatggtgagc
aagggtgcggagctgttcaccggtatcgtgccgatcctgattgaactgaacggcgacgtgaacggtcacaagttcagcgtgagcggcgagggtgaag
gcgatgccacctacggcaagctgaccctgaaattcatttgcaccaccggcaagctgccagtgccatggccaaccctggtgaccaccctgagctacggtg
tgcagtgctttagccgttatccggaccacatgaagcagcacgatttctttaaaagcgccatgccggagggctacatccaggaacgtaccattttctttgagg
acgatggtaactataagagccgcgccgaggtgaaattcgaaggcgacaccctggtgaaccgtatcgagctgaccggtaccgactttaaggaagatggc
aacattctgggtaacaaaatggagtacaactataacgcgcacaacgtgtacatcatgaccgataaggctaaaaacggcattaaggtgaacttcaaaatcc
gccacaacattgaagacggtagcgtgcagctggcggatcactaccagcagaacaccccgatcggtgacggtccagtgctgctgccggataaccactat
ctgagcacccagagcgccctgagcaaggacccgaacgagaaacgtgatcacatgatctacttcggctttgtgaccgccgcggctattacccacggtatg
gacgagctgtacaagaccctgggcatggatgaactgtataaaggtagcggcggtagcggcggtagcggcggtagcggcggtagcggcggtagcgg

ggctgaggccgcggctaaggaagccgcggctaaagaggccgcggctaaggaagccgcggctaaagaggccgcggctaaggaagccgcggctaa
ggctggcagcggcggtagcggcggtagcggcggtagcggcggtagcggcggtagcggtgccgaagccgcggctaaggaagccgcggctaaaga
ggccgcggctaaggaagccgcggctaaagaggccgcggctaaggaagccgcggctaaagccggcagcggcggtagcggcggtagcggcggtag
cggcggtagcggcggtagcggcgtgagcaagggcgagatggtgagcaaaggcgaggaagacaacatggccatcattaaggagttcatgcgctttaa
agtgcacatggaaggcagcgtgaacggtcacgagtttgaaatcgagggtgaaggcgagggtcgtccgtacgagggtacccagaccgctaagctgaaa
gtgaccaagggcggtccactgccattcgcttgggacatcctgagcccgcagtttatgtacggcagcaaggcctatgtgaaacacccggcggacattccg
gattatctgaagctgagcttcccggagggttttaaatgggaacgcgtgatgaacttcgaggatggcggtgtggtgaccgtgacccaggacagcagcctg
caggatggcgaattcatctacaaggtgaaactgcgtggcaccaactttccgagcgacggtccggttatgcagaagaaaaccatgggttgggaggccag
cagcgaacgcatgtatccggaggatggcgcgctgaagggcgagatcaagcagcgtctgaagctgaaagacggcggtcactacgatgctgaggtgaa
gaccacctataaagccaagaaaccggtgcagctgccgggcgcttacaacgtgaacatcaagctggacattaccagccacaacgaggattacaccatcg
tggaacagtatgagcgtgctgaaggtcgtcacagcaccggcggtatggacgaactgtataaa

Color legends for amino acid and nucleic acid sequences: N-terminal tags, AcGFP1, flexible linker with $(GSG)_6$, a rigid α-helical region with $(EAAAK)_6$ repeats, and mCherry

**Characterization of CrH2, before and after ultracentrifugation**

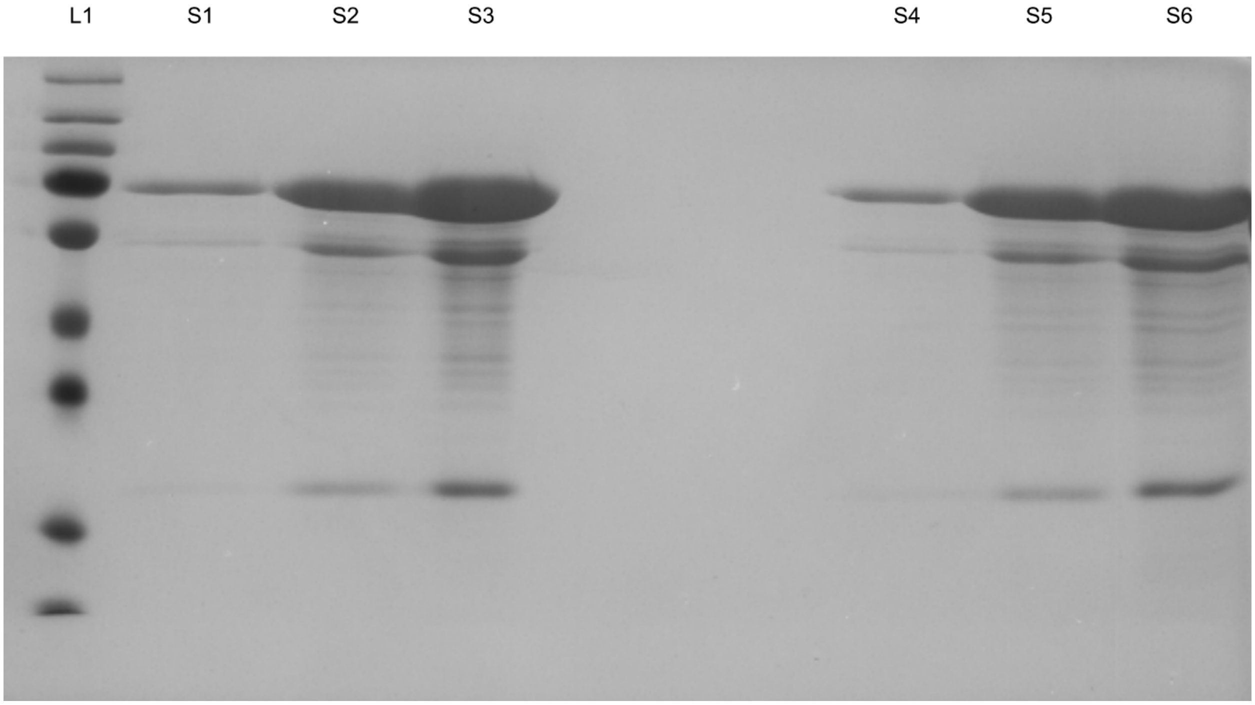


*Figure S1. SDS-PAGE of CrH2 before and after ultracentrifugation. Lane L1: 5 μL of protein ladder (Promega #26619), S1-S3: post-ultracentrifugation samples at 1, 5, and 10 μM (20 μL each), and S4-S6: pre-ultracentrifugation samples at 1, 5, and 10 μM (20 μL each), respectively. We do not see significant protein loss before or after ultracentrifugation.*

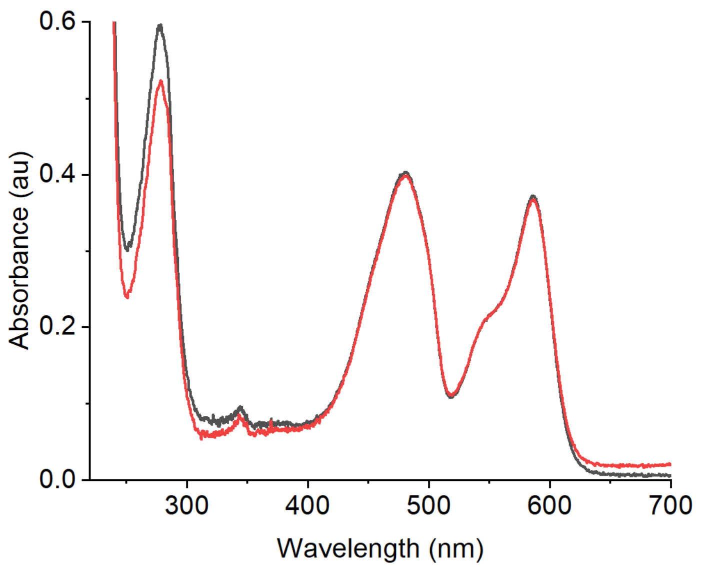


*Figure S2. The UV/Vis spectra of 5 µM CrH2 in 20 mM HEPES and 150 mM NaCl, pH 7.5, are black before ultracentrifugation and red after ultracentrifugation.*

### Computed physical parameters of CrH2

*Table S1. Extinction coefficients of AcGFP1, mCherry and $\varepsilon_{280}$ (CrH2).*

| | $\varepsilon(M^{-1}cm^{-1})$ | Wavelength (nm) |
|---|---|---|
| mCherry[3] | 72,000 | 587 nm |
| AcGFP1[3] | 32,500 | 475 nm |
| $\varepsilon_{280}$ (CrH2)[4] | 60,740* | 280 nm |

*The extinction coefficients of CrH2 at 280 nm were calculated from the amino acid sequence using Expasy's Protparam tool (https://web.expasy.org/protparam).[4]

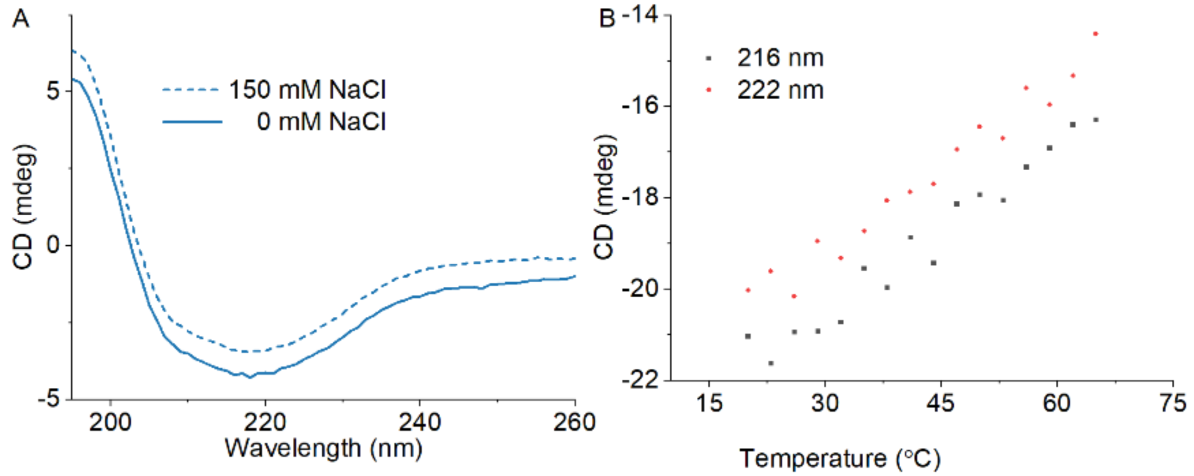


*Figure S3. CD spectra of CrH2 in 20 mM phosphate buffer, pH 7.0. A) The CD spectra of CrH2 remain qualitatively similar with or without 150 mM NaCl. B) The CD thermal melt of 2.5 µM CrH2 at 216 and 218 nm was acquired in a Jasco spectrofluorometer FP-8300 in 20 mM phosphate buffer, pH 7.0, from 20°C up to 65°C in steps of 3°C in a 1 mm quartz cuvette. A steady change in the CD, unlike a typical sigmoidal thermal denaturation for folded proteins,[5] represents stability and uncooperative secondary structure transitions during thermal melting.[1] However, a slight decrease in the CD at 222 nm with temperature perturbation may represent a decrease in the helical content.[1]*

### Spectra comparison of CrD with Cy3/Cy5 DNA sensor

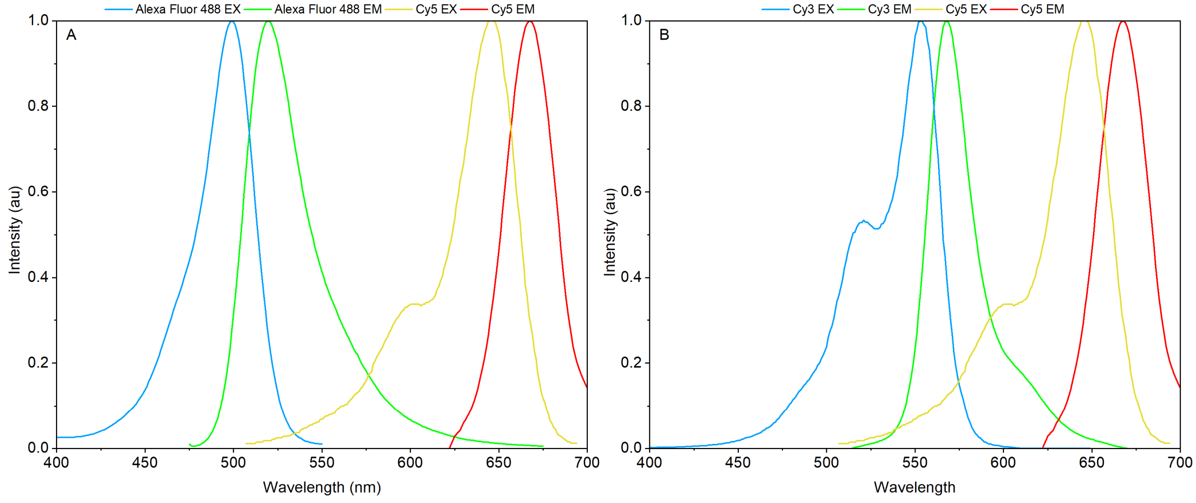


*Figure S4. Spectral comparison of Alexa488/Cy5-containing CrD (Table 1) vs. Cy3/Cy5-based DNA sensor (Shubeita, G. Personal Email Correspondence. May 16, 2023).[6] The spectra were simulated using the FPbase website (https://www.fpbase.org/spectra/).*

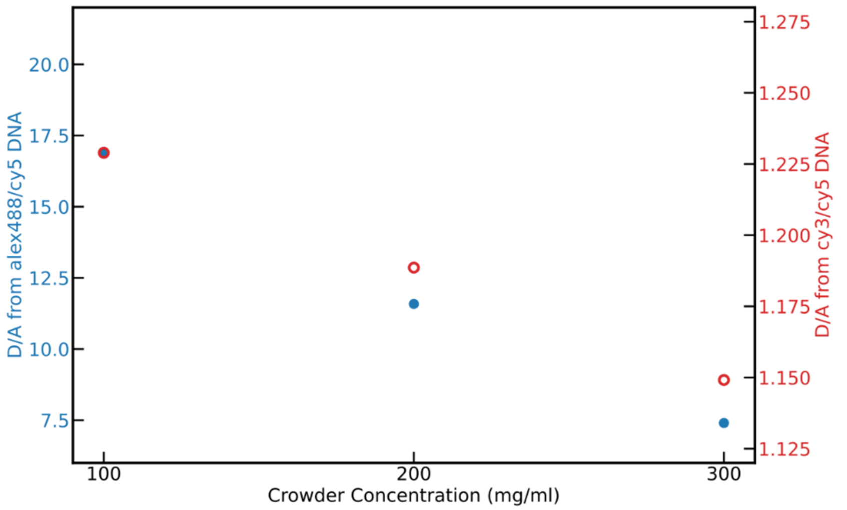


*Figure S5. The fluorimetric FRET-based comparison of the crowding response of CrD (blue filled circles) vs Cy3/Cy5-based DNA sensor (red empty circles) with 100 – 300 mg/mL of PEG 8 kDa. The left and right-hand side axes represent the donor/acceptor-FRET ratios of the DNA-based crowding sensors.*

*Table S2. Comparison of the DNA sensor with Alexa488 vs. Cy3 as a donor fluorophore.[7,8]*

| FRET pair | **QY**$_{Donor}$ | **EC**$_{Acc.}$ ($M^{-1}$ $cm^{-1}$) | **QY**$_{Acc.}$ | $J(\lambda)$ (*1e15 $M^{-1}$ $cm^{-1}$ $nm^{4}$) | $R_0$(Å) | $R_0 \times QY_A$ | $\kappa^2$ | $n$ |
|---|---|---|---|---|---|---|---|---|
| Alexa488/Cy5 | 0.92 | 250,000 | 0.3 | 1.83 | 56.21 | 16.86 | 0.6667 | 1.33 |
| Cy3/Cy5 | 0.15 | 250,000 | 0.3 | 6.32 | 51.09 | 15.33 | 0.6667 | 1.33 |

QY = Quantum Yield, EC = Extinction Coefficient, J(λ) = Overlap Integral, R0 = Förster radius, $n$ = refractive index, $\kappa^2$ = orientation factor.[8] The equation 1 and 2, provided below, calculates $R_0$ and J(λ).[8]

$$R_0 = 0.211. \sqrt[6]{k^2 n^{-4} Q_D J(\lambda)} \textit{Equation 1)}$$

$$J(\lambda) = \frac{\int F_D(\lambda)\epsilon_A(\lambda)\lambda^4 d\lambda}{\int F_D(\lambda)\, d\lambda} \textit{Equation 2)}$$

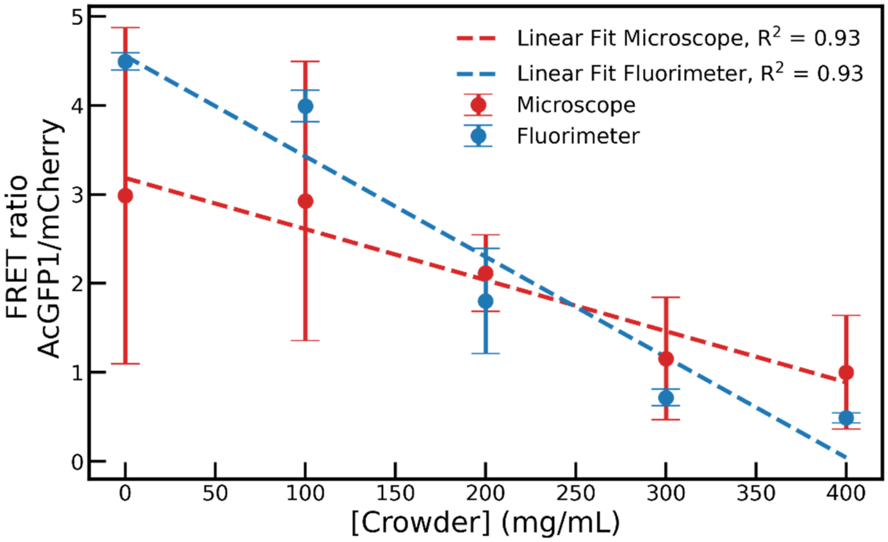


*Figure S6. A linear regression fit of the CrH2 crowding sensor response (Figure 1H) from the fluorimeter (blue color) and the microscope (red color). The crowder refers to PEG 8 kDa at 0–400 mg/mL.*

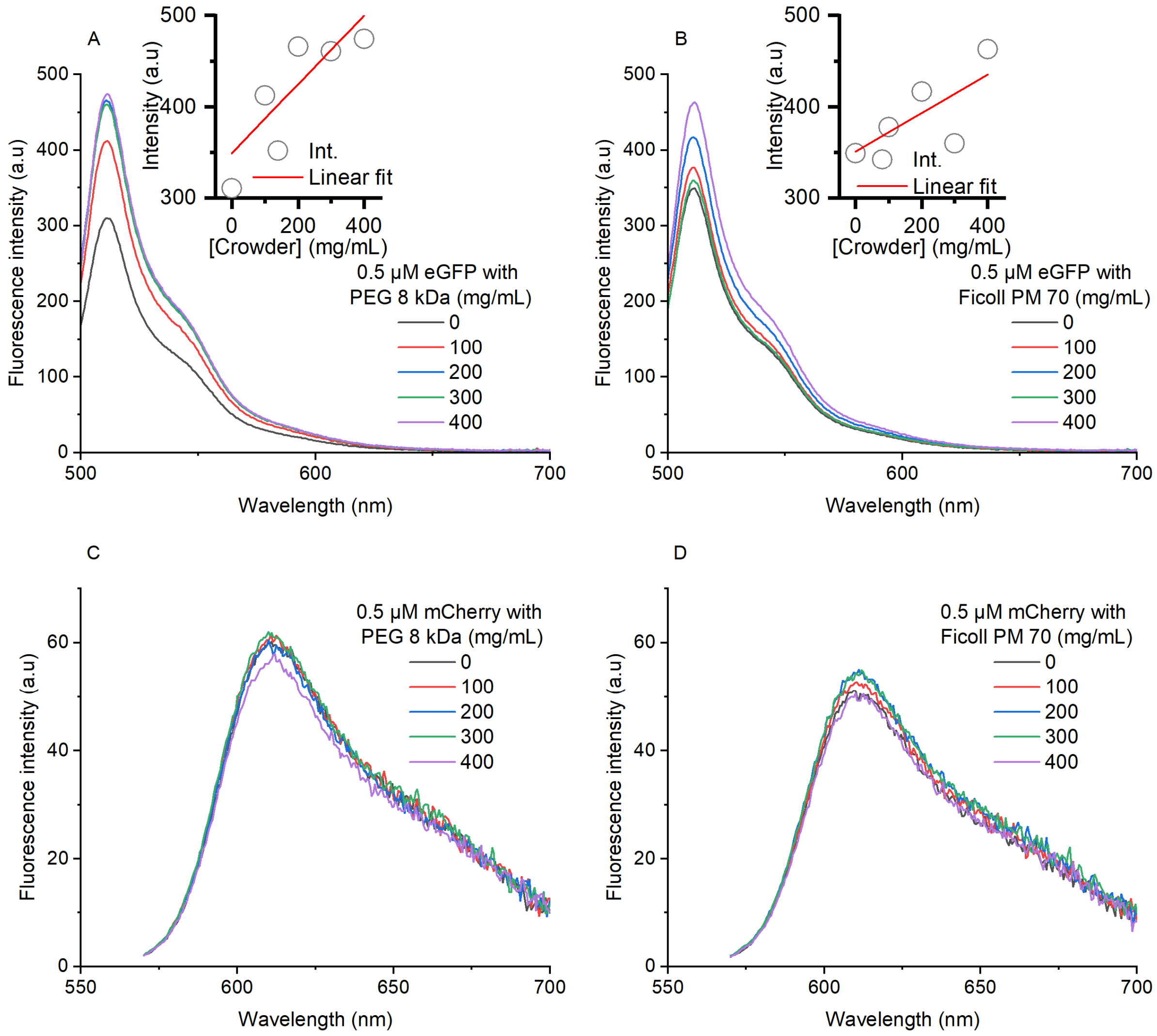


*Figure S7. Fluorescence emission spectra of eGFP and mCherry in crowding conditions. (A and B) Emission spectra of 0.5 µM of eGFP in the presence of 0 – 400 mg/mL of A) PEG 8 kDa and B) Ficoll PM 70. The insets show an intensity linear regression plot. We see an emission enhancement of eGFP in the presence of PEG and Ficoll. (C and D) Emission spectra of 0.5 µM of mCherry in the presence of 0 – 400 mg/mL of C) PEG 8 kDa and D) Ficoll PM 70. We don't observe a significant change in the emission spectra of mCherry in the presence of PEG and Ficoll. The buffer solution contains 20 mM HEPES, 150 mM NaCl, pH 7.5, 1 mM Trolox, and 1 mM DTT. Excitation at 488 nm and 561 nm, and emission from 500 – 700 nm and 570 – 700 nm, for eGFP and mCherry, respectively. Excitation and emission bandwidths are set at 2.5 nm, with a 0.1 nm response time, 0.5 nm data interval, a 500 nm/min scan speed, and a 25°C cell temperature. The samples were repeated twice, and a representative spectrum is presented.*

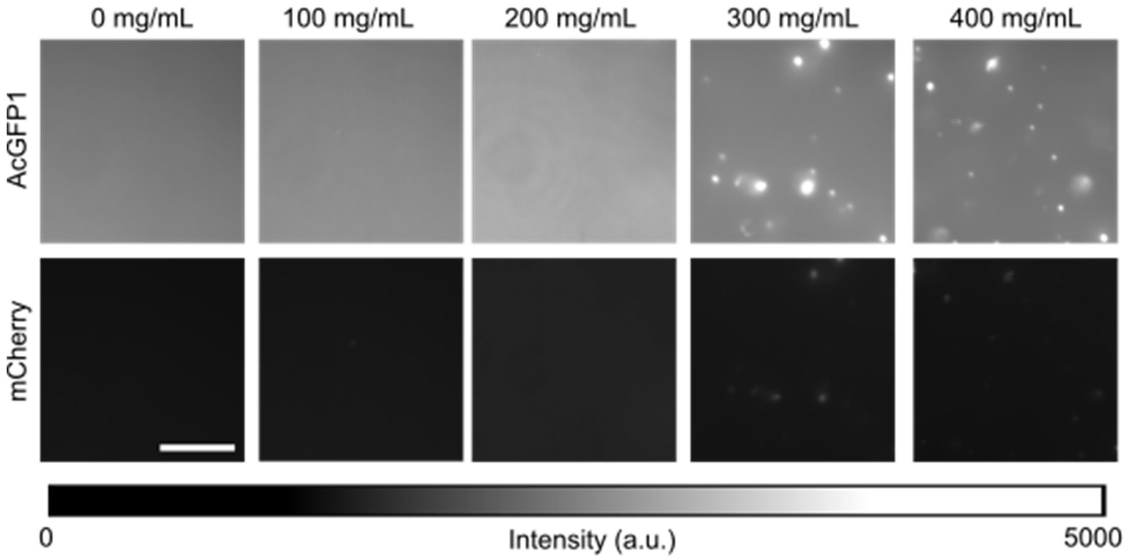


*Figure S8.* *A 2 µM CrH2 sample phase separates in the presence of 200 – 400 mg/mL of PEG 8 kDa at minimal salt concentrations. Buffer: 20 mM sodium phosphate buffer, pH 7.5, without any additional NaCl was used.*

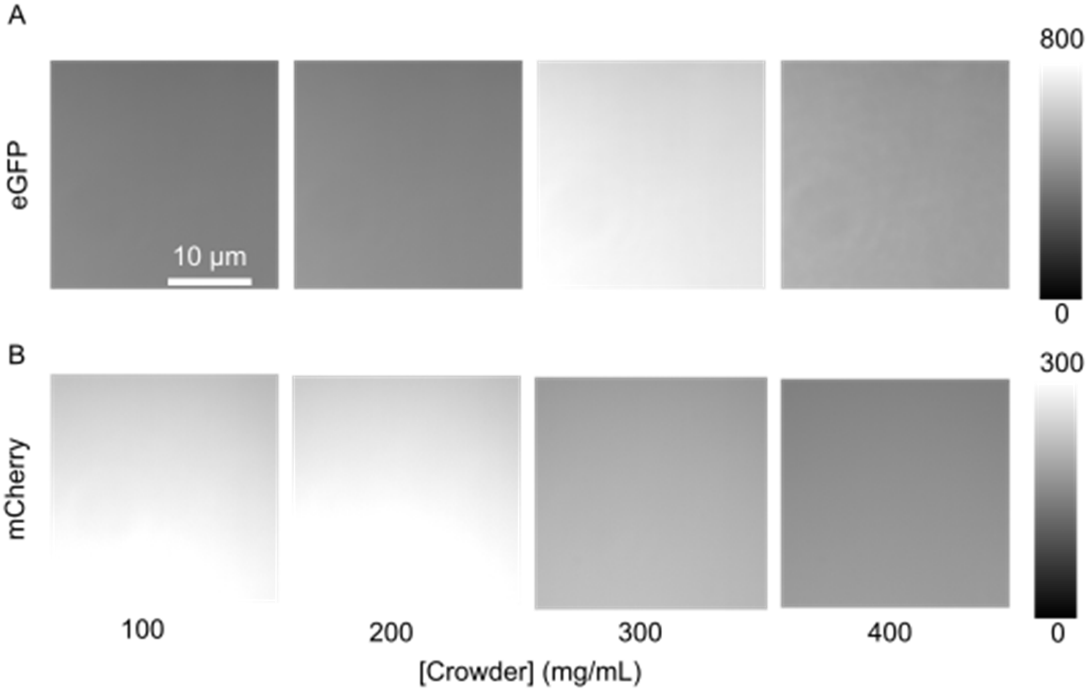


*Figure S9. The eGFP and mCherry control samples do not phase separate in the presence of PEG 8 kDa. Imaging of 0.5 µM A) eGFP using 488 nm laser or B) mCherry with 561 nm laser. The individual samples contain 100–400 mg/mL PEG in 20 mM HEPES, 150 mM NaCl, pH 7.5, 1 mM DTT, and 1 mM Trolox buffer.*

*Table S3. Calculation of excluded volume fractions*

| | | Ficoll PM 70 | | | PEG 1 kDa | | | PEG 8 kDa | | | PEG 20 kDa | | | PEG 35 kDa | |
|---|---|---|---|---|---|---|---|---|---|---|---|---|---|---|---|
| Crowder, % | Excluded volume fraction (v/v) | D/A | SD | Excluded volume fraction (v/v) | D/A | SD | Excluded volume fraction (v/v) | D/A | SD | Excluded volume fraction (v/v) | D/A | SD | Excluded volume fraction (v/v) | D/A | SD |
| 0 | 0.00 | 4.50 | 0.16 | 0.00 | 4.56 | 0.14 | 0.00 | 4.43 | 0.00 | 0.00 | 4.42 | 0.02 | 0.00 | 4.16 | 0.44 |
| 10 | 0.48 | 4.34 | 0.22 | 0.11 | 4.33 | 0.15 | 0.38 | 3.87 | 0.03 | 0.52 | 3.42 | 0.30 | 2.06 | 4.11 | 0.45 |
| 20 | 0.96 | 4.26 | 0.16 | 0.21 | 3.90 | 0.22 | 0.76 | 1.39 | 0.11 | 1.04 | 0.91 | 0.01 | 4.12 | 1.01 | 0.04 |
| 30 | 1.43 | 4.08 | 0.19 | 0.32 | 3.23 | 0.47 | 1.14 | 0.65 | 0.02 | 1.55 | 0.56 | 0.00 | 6.19 | 0.57 | 0.00 |
| 40 | 1.91 | 3.92 | 0.17 | 0.43 | 1.10 | 0.18 | 1.51 | 0.46 | 0.01 | 2.07 | 0.39 | 0.01 | 8.25 | 0.44 | 0.00 |

Note: Refer to Table S4 below for total excluded volume calculations.

*Table S4. Calculation of total excluded volume from hydrodynamic radius*

The hydrodynamic radius and excluded volume occupied by polymers in the solution state vary with solution conditions and measurement approaches. To simplify and approximate the excluded volume fraction contributed by PEG and Ficoll, we assumed the polymer particles to be hard spheres. We have calculated the volume occupied by the polymer using the hydrodynamic radius determined experimentally elsewhere.[9–13]

| Crowder | MW (g/mol) | Rh (nm) | $Rh^3$ ($nm^3$) | Volume of 1 particle ($4/3\pi Rh^3$) ($nm^3$) | Volume of 1 particle ($4/3\pi Rh^3$) ($cm^3$) | Moles in 10 g of crowder solution | Number of Particles (N) in 10 g | Vev, Total Excluded Volume (mL) in 100 mL of 10 % (w/v) crowder | Vev, Total Excluded Volume (mL) in 100 mL of 20 % (w/v) crowder | Vev, Total Excluded Volume (mL) in 100 mL of 30 % (w/v) crowder | Vev, Total Excluded Volume (mL) in 100 mL of 40 % (w/v) crowder |
|---|---|---|---|---|---|---|---|---|---|---|---|
| Ficoll PM70 | 70000 | 5.1[9] | 132.7 | 555.4 | 5.55E-19 | 1.43E-04 | 8.60E+19 | 47.8 | 95.6 | 143.4 | 191.1 |
| PEG 1 kDa | 1000 | 0.75[13] | 0.4 | 1.8 | 1.77E-21 | 1.00E-02 | 6.02E+21 | 10.6 | 21.3 | 31.9 | 42.6 |
| PEG 8 kDa | 8000 | 2.29[12] | 12.0 | 50.3 | 5.03E-20 | 1.25E-03 | 7.53E+20 | 37.9 | 75.7 | 113.6 | 151.4 |
| PEG 20 kDa | 20000 | 3.45[10] | 41.1 | 171.9 | 1.72E-19 | 5.00E-04 | 3.01E+20 | 51.8 | 103.5 | 155.3 | 207.1 |
| PEG 35 kDa | 35000 | 6.59[11] | 286.2 | 1198.2 | 1.20E-18 | 2.86E-04 | 1.72E+20 | 206.2 | 412.4 | 618.6 | 824.8 |

Note: $V_{ev}$ is the total excluded volume by the crowder, $V_t$ is the total volume of the solution i.e. 100 mL, excluded volume fraction = $V_{ev}/V_t$, Avogadro's Number (NA) = $6.022\times10^{23} mol^{-1}$; Volume of a sphere = $4/3\pi Rh^3$; Conversion: 1 $nm^3 = 10^{-21}$ $cm^3$ or $10^{-21}$ mL; A 10% w/v concentration means 10 grams of PEG in 100 mL of solution.

**The sequence properties potentially drive the phase separation of CrH2**

To gain insights into the phase behavior of the CrH2 protein polymer, we deep-dived into its sequence composition.[14,15] The CrH2 has two globular fluorophore domains connected through linkers and helical polypeptide stretches.[1,16] The linker polypeptide with a sequence of "TLGMDELYK $(GSG)_6A(EAAAK)_6(GSG)_6A(EAAAK)^6A(GSG)_6$VSKGE" houses two helical peptide stretches, designated as "$(EAAAK)_6$", and flexible linker motifs, defined by "GSG". The repeating stretches of Ser (S) and Gly (G) residues in the linker region are likely to have a "sticker"-like role in guiding the phase separation of CrH2.[17,18] Moreover, the purification procedures reported elsewhere render an IDR-like additional poly-histidine tag at the N-terminus of CrH2, adding to the cumulative effects from small stretches of IDRs in the linker polypeptide.[1,16]

We have used FuzDrop[19–21] to understand the droplet-promoting propensity of CrH2 (Figure S10). The server predicts a pLLPS score based on the free states' conformational entropy and the protein's binding entropy. A higher pLLPS score of 0.9 from FuzDrop suggests that the protein will likely undergo phase separation. Thus, PEG, a known inducer of phase separation, may likely enhance the inherent tendency of CrH2 to undergo phase separation. Although we observe phase separation for CrH2 in the presence of PEG (Figure 2), we do not see similar behavior with the eGFP or mCherry fluorophore alone samples under identical conditions (Figure S9). This drives us to emphasize the importance of the linker region in driving the phase separation of CrH2.

The linker framework used in CrH2 by Boersma et al. was inspired by the work of Golynskiy et al. on a cerulean/citrine FRET-based HIV-1 antibody detection sensor.[22,23] The earlier probe detects bivalent antibodies by binding to the Fab arms 120–170 Å apart. The authors used the cerulean and citrine domains as FRET pairs, respectively.[23] They included critical hydrophobic mutations (S208F/V224L and Q204F/V224L), which rendered the intra-domain stickiness, resulting in the enhanced emission response of the probe in the off state. Further, the conformationally flexible linker, flanking the two fluorophore domains, carries two rigid alpha helices and two HIV-1 epitope sequences, "EKIRLR". The alpha helices were inspired by the work of Marqusee et al., where they discovered a slight stretch of peptide sequence adopting alpha-helical conformation at low temperatures and salt concentrations.[24] Poolman and co-workers repurposed the cerulean/citrine probe to sense macromolecular crowding. The probe carries the A206K mutation and lacks the hydrophobic "sticky" mutations in both fluorophore domains to minimize self-association and, therefore, could obtain a high donor/acceptor FRET ratio in the absence of crowding.[23,25] The HIV-1 epitope sequences in the linker region were removed, leaving behind the current linker of the CrH2.

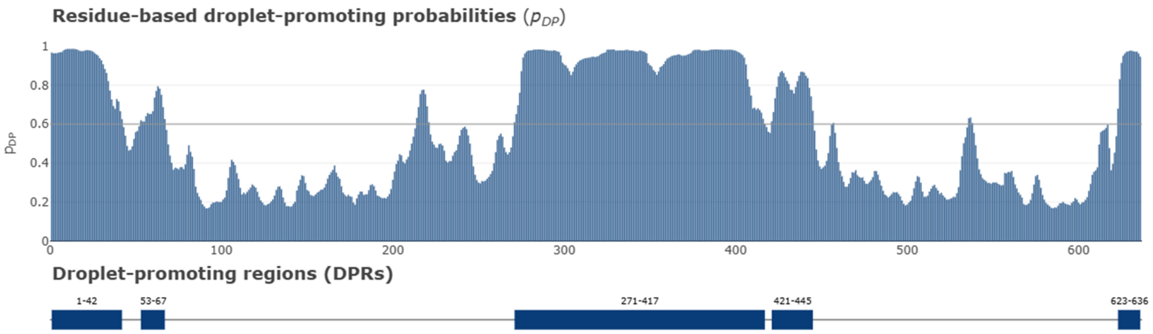


*Figure S10. Prediction of droplet-promoting propensity of CrH2 using FuzDrop, with pLLPS = 0.8900.*
